# Sequence-encoded autoinhibition couples mRNA decapping activity to phase separation

**DOI:** 10.64898/2026.06.16.732745

**Authors:** Trase Aguigam, Katarzyna Grab, Joanna Kowalska, Jacek Jemielity, John D. Gross

## Abstract

Removal of the 5′ m⁷G cap by the Dcp1/Dcp2 complex commits mRNAs to degradation, yet the mechanisms regulating decapping remain incompletely understood. Here, we identify residue-level determinants within the extended C-terminus of fission yeast Dcp2 that repress activity. Mutations in conserved inhibitory motifs relieve autoinhibition, enhance RNA binding, and bypass the requirement for the activator Edc3. Strikingly, this activation persists within phase-separated condensates, demonstrating that conformational relief in solution is propagated to the dense phase. We further show that long-range interactions between the intrinsically disordered region of Dcp2 and the catalytic core restrict RNA engagement, providing a mechanistic basis for negative regulation. Together, these findings establish that sequence-encoded elements within the Dcp2 C-terminus control catalytic activity and functional output within biomolecular condensates. More broadly, our results reveal that competing interactions encoded within intrinsically disordered regions of proteins are balanced to allosterically tune enzyme activity, providing a general mechanism by which proteins modulate distinct enzymatic functions within biological condensates.

## Introduction

The 5’ N7-methylguanosine (m^7^G) cap is a defining feature of the 5’ end of eukaryotic mRNAs and is involved in the regulation of gene expression, nuclear mRNA export, cap-dependent translation initiation, and mRNA quality control (Moore 2005; Isken and Maquat 2007; Aregger and Cowling 2016; Ramanathan et al. 2016). Removal of the 5’ cap (decapping) is a conserved cellular process from yeast to humans. It represents a key step in the lifecycle of cytoplasmic mRNAs, committing transcripts to 5’-3’ degradation by the exoribonuclease Xrn1, and thereby shaping gene expression profiles (van Hoof and Parker 2002; Coller and Parker 2004; Dowdle and Lykke-Andersen 2025). In eukaryotes, decapping is catalyzed by the conserved Dcp1/Dcp2 decapping complex, a bipartite holoenzyme whose activity is tightly regulated to ensure appropriate temporal and spatial control of mRNA decay (Coller and Parker 2004; Parker and Sheth 2007) (Fig. 1a). Structural studies of the Dcp2 catalytic and regulatory domains reveal a dynamic composite active site that is occluded or accessible depending on the presence of cap analogs and protein cofactors (She et al. 2006; Floor et al. 2010; Mugridge et al. 2016; Charenton et al. 2016; Wurm et al. 2017; Mugridge et al. 2018b). However, it has become increasingly clear that decapping regulation extends beyond the structured core domains. In particular, the extended C-terminal intrinsically disordered region (IDR) of yeast Dcp2 has been shown to play a central role in modulating decapping activity through protein–protein interactions and context-dependent regulatory inputs (Jonas and Izaurralde 2013; He and Jacobson 2015; Paquette et al. 2018; He et al. 2018, 2022; He and Jacobson 2023).

**Figure 1.**
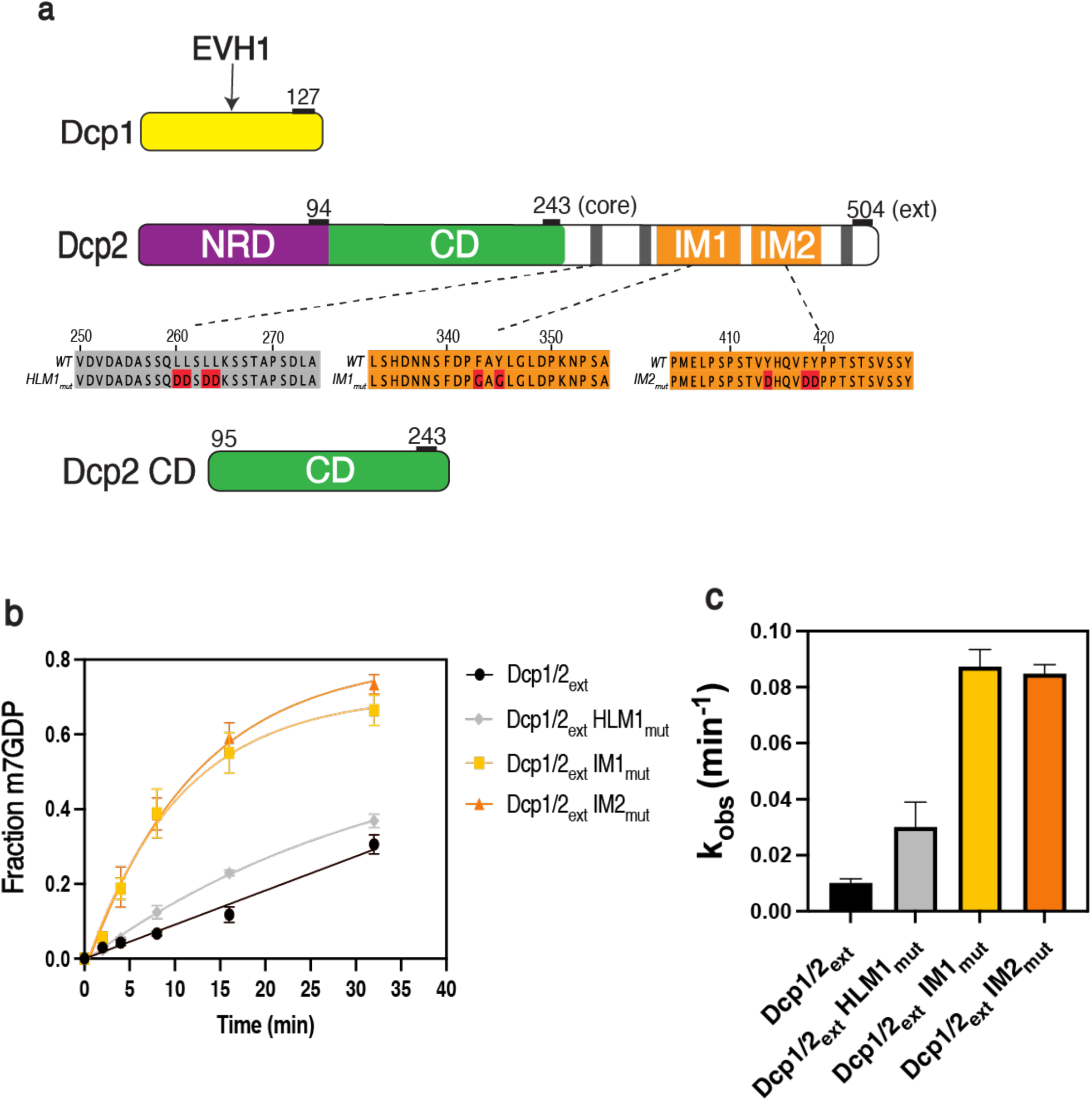
The C-terminal extension of Dcp2 confers autoinhibition. **a,** Schematic domain organization of Dcp1, full-length Dcp2 extension construct (Dcp2_ext_) and the isolated catalytic core (Dcp2 CD) used in this study. Dcp1 contains an EVH1 domain. Dcp2 comprises an N-terminal regulatory domain (NRD), catalytic domain (CD), and a C-terminal extension harboring helically leucine-rich motifs (HLMs) and inhibitory motifs (IM1 and IM2). Mutated residues analyzed in this study are indicated. **b,** Representative time course of m^7^GDP production comparing wild-type Dcp1/Dcp2_ext_ with mutants disrupting IM1, IM2, or HLM1. Data are plotted as the fraction of product formed over time. **c,** Observed rate constants (k_obs_) derived from fits to the progress curves shown in panel **b**.

Within the C-terminal IDR of yeast Dcp2, short linear interaction motifs (SLiMs) function as both inhibitory and activating regulatory elements (He and Jacobson 2015; Paquette et al. 2018; He and Jacobson 2023). Dcp1 is an obligate cofactor and protein interaction platform that binds proline-rich motifs found in activators of decapping or inhibitory motifs (IMs) within the Dcp2 C-terminus (She et al. 2004; Borja et al. 2011; Lai et al. 2012; Wurm et al. 2016; Mugridge et al. 2018a) (Figure 1a). Dcp1 and its interactions with IMs are thought to stabilize an inactive conformation of the Dcp1/Dcp2 complex, limiting access to the RNA-binding channel and suppressing catalytic activity (Wurm et al. 2016, 2017; Paquette et al. 2018). These findings have led to a model in which IMs are spatially positioned near the structured core domains, reinforcing a closed, inactive conformation incompatible with substrate binding and catalysis (Paquette et al. 2018). This regulatory strategy is reminiscent of autoinhibitory mechanisms observed in other systems, such as the eukaryotic proto-oncogene protein kinase Src (Sicheri et al. 1997; Huse and Kuriyan 2002; Pufall and Graves 2002; Davey et al. 2011). The binding of the enhancer of decapping protein 3 (Edc3) to adjacent helical leucine-rich motifs (HLMs) was proposed to relieve autoinhibition by liberating IMs from the Dcp2 core domains and promoting an open precatalytic conformation of the enzyme that can bind capped RNA and subsequently form a closed, substrate-bound conformation with residues in the composite active site poised for catalysis (Fromm et al. 2012; He and Jacobson 2015; Charenton et al. 2016; Wurm et al. 2017; Paquette et al. 2018). Together, these observations support a model in which the Dcp2 C-terminus functions as a modular regulatory platform that integrates inhibitory and activating cues to exert conformational control over decapping (Mugridge et al. 2018a, 2018b). However, most prior studies have relied on large truncations or deletions of the IDR, limiting insight into the contributions of individual residues and precluding a residue-level understanding of how specific sequence motifs regulate autoinhibition, substrate recognition, and enzyme activation (He and Jacobson 2015; Paquette et al. 2018).

Recent work on biological condensates has introduced an additional layer of complexity to Dcp1/Dcp2 regulation (Banani et al. 2017; Schütz et al. 2017; Tibble et al. 2021). It has been proposed that biological condensates represent non-canonical modes of cellular organization, coupling reaction directionality with concentration-dependent effects to accelerate enzymatic reactions (Ditlev et al. 2018; Peeples and Rosen 2021; O’Flynn and Mittag 2021). Consistent with this idea, Dcp1/Dcp2 and its coactivators, including Edc3, are enriched in processing bodies (P-bodies), phase-separated cytoplasmic condensates whose assembly is driven by multivalent interactions mediated by IDRs and short linear motifs (Sheth and Parker 2003; Parker and Sheth 2007; Jonas and Izaurralde 2013; Harmon et al. 2017; Xing et al. 2020; Currie et al. 2022). Notably, the Dcp2 IDR is necessary for entry into P-bodies in vivo and drives LLPS in vitro (Harigaya et al. 2010; Fromm et al. 2014; Xing et al. 2020). In this framework, enrichment within P-bodies has been proposed to promote decapping by increasing encounter frequency between reactants and stabilizing productive molecular interactions, particularly for low-affinity or diffusion-limited reactions (Decker and Parker 2012; Fromm et al. 2014; Luo et al. 2018; Klosin et al. 2020; Riback et al. 2020). Additionally, HLMs within the Dcp2 IDR support multivalent binding to the Lsm domain of Edc3, rewiring protein-protein interactions to drive LLPS, and forming catalytically active condensates (Fromm et al. 2012; Tibble et al. 2021). However, emerging evidence also suggests that inhibitory motifs may influence phase separation and activity within condensates by reinforcing autoinhibitory interactions between the IDR and the structured core domains (Tibble et al. 2021). These seemingly opposing effects of IDR mediated interactions imply a complex interplay between distinct interfaces. Dissecting this interplay requires an understanding of the specific sequence determinants governing condensate organization, protein partitioning, and the functional coupling between autoinhibition and phase separation. However, such determinants remain incompletely understood.

Here, we dissect the molecular contributions of individual sequence motifs within the Dcp2 IDR to autoinhibition, RNA binding, and condensate organization using a combination of targeted mutagenesis, quantitative biochemical assays, and in vitro reconstitution of phase-separated biological condensates. We show that IM-mediated autoinhibition directly suppresses RNA engagement and that targeted disruption of these motifs enhances substrate binding and bypasses the requirement for Edc3 in decapping activation, directly testing the role of an IDR-stabilized closed conformation. Furthermore, we demonstrate that the interactions responsible for autoinhibition in solution are also required for autoinhibition within phase-separated condensates. Specific lesions within IM2 stimulate decapping within reconstituted Dcp1/Dcp2 condensates in the absence of coactivators, revealing that sequence grammar within the Dcp2 IDR organizes the multivalent interaction networks that underlie phase separation.

Together, our findings establish a residue-level framework for understanding how the Dcp2 IDR integrates inhibitory and activating signals to control mRNA decapping. Importantly our studies suggest more broadly that condensates can stabilize one of several transient conformational states that exist in dilute phase to regulate enzyme activity through a ‘conformational selection’ mechanism.

## Results

### Aromatic residues within the C-terminal IDR of Dcp2 repress decapping

To investigate the contribution of specific sequence motifs within the Dcp2 IDR, we co-expressed Dcp1 with a Dcp2 construct containing the IDR up to residue 504 (Dcp1/Dcp2_ext_) as previously described (Paquette et al. 2018; Tibble et al. 2021) (Fig. 1a). This construct was selected based on its optimal expression while retaining the regulatory elements necessary for autoinhibition and activation. Dcp1/Dcp2_ext_ contains an N-terminal regulatory domain (NRD), a catalytic domain (CD), and a C-terminal IDR containing three HLMs and two IMs. Because prior models of Dcp1/Dcp2 regulation have relied largely on internal deletions of the Dcp2 IDR, we instead employed targeted point mutations to perturb discrete sequence features without altering overall IDR length or domain architecture.

We focused on conserved aromatic residues within IM1 and IM2 and conserved leucine residues within HLM1 (Fig. 1a). These residues were selected for mutational analyses due to their conservation across multiple *Schizosaccharomyces* species, suggesting functional importance (Supplementary Figure S1).

To assess the molecular determinants of autoinhibition by the Dcp2 IDR, we compared the decapping activity of wild-type Dcp1/Dcp2_ext_ with that of complexes bearing point mutations in conserved residues of IM1 or IM2 (IM1_mut_, IM2_mut_, respectively) (Fig.1a). We also introduced a mutation in HLM1 (HLM1_mut_) to investigate how this interaction motif affects decapping activity, since prior structural and biochemical studies suggested it may promote the autoinhibited conformation of Dcp1/Dcp2 (Fromm et al. 2012; Mugridge et al. 2018a). Decapping reactions were performed using cap-radiolabeled 29-mer RNA substrates as described previously (She et al. 2006; Jones et al. 2008; Paquette et al. 2018). All reactions were conducted under sub-saturating, single-turnover conditions to minimize effects arising from phase separation, multivalent interactions, or enzyme recycling, enabling accurate measurement of mRNA decapping rates that include effects from binding and catalysis (k_obs_; see Methods). Mutation of conserved aromatic residues in either IM1_mut_ or IM2_mut_ resulted in an approximately tenfold increase in decapping activity relative to wild type. In contrast, HLM1_mut_ produced only a modest enhancement (Fig. 1b, c). Importantly, the observed increases in enzymatic activity were not attributable to variations in sample preparation or oligomerization. All decapping complexes eluted as a single, well-resolved and homogeneous peak by gel-filtration chromatography and did not exhibit detectable oligomerization under the decapping reaction conditions tested (Supplementary Figure S2).

These data support a model in which specific aromatic residues within IM1 and IM2 stabilize inhibitory intramolecular contacts that suppress activity. The limited impact of HLM1 is consistent with a model in which HLMs primarily mediate intermolecular interactions with activating cofactors such as Edc3, rather than directly regulating access to the active site. Taken together, these enzymatic data reveal a functional partitioning within the Dcp2 IDR, with IMs driving intramolecular autoinhibition and HLM1 contributing modestly to this effect.

### Multiple IMs contribute to the suppression of RNA binding

The effects observed above could arise from effects on binding and catalysis. To dissect the contribution to each step, we next investigated how mutations within sequence motifs of the Dcp2 IDR influence the ability of the decapping complex to bind RNA. Structural and biochemical studies have shown that Dcp2 contains a positively charged RNA-binding channel that spans the junction between the catalytic and regulatory domains and extends toward the Dcp1-binding interface, forming an electrostatically favorable surface that engages the RNA body and positions the 5′ mRNA cap into the composite active site for hydrolysis (Mugridge et al. 2018a). Given that low-complexity regions can form transient contacts with folded domains (Pufall and Graves 2002; Tompa 2012; Dyson and Wright 2005; Borgia et al. 2018), we hypothesized that IMs may be positioned near the structured core to reinforce autoinhibition by partially occluding or destabilizing access to the RNA-binding channel.

To test this hypothesis, we quantified RNA binding across a series of Dcp1/Dcp2 constructs differing in IDR composition using fluorescence polarization (FP). FP measurements were performed using a 5′-phosphorylated poly-U30 RNA oligonucleotide with a 3′-6-FAM modification, which serves as a minimal substrate mimic. In this assay, the polarization signal increases proportionally with substrate binding, enabling precise determination of the dissociation constant (K_d_; see Methods) (Moerke 2009). As a positive control, we used a Dcp2 construct consisting only of the catalytic domain (Dcp2 CD), which has previously been reported to have the highest RNA-binding affinity (Wurm et al. 2017; Tibble et al. 2021) (Fig. 1a).

Consistent with prior reports, Dcp2 CD bound RNA with low-micromolar affinity (Wurm et al. 2017; Tibble et al. 2021) (Fig. 2a, b). In contrast, wild-type Dcp1/Dcp2_ext_ exhibited substantially weaker RNA binding, consistent with a role for the IDR in reinforcing autoinhibition. Notably, IM1_mut_ and IM2_mut_ enhanced RNA binding by approximately tenfold relative to wild-type, restoring affinities comparable to those of IDR-truncated or core-only constructs (Dcp2_core_). These data indicate that IM1 and IM2 function to restrict accessibility to the RNA-binding channel, likely by stabilizing an autoinhibited conformation through transient intramolecular contacts with the structured core.

**Figure 2.**
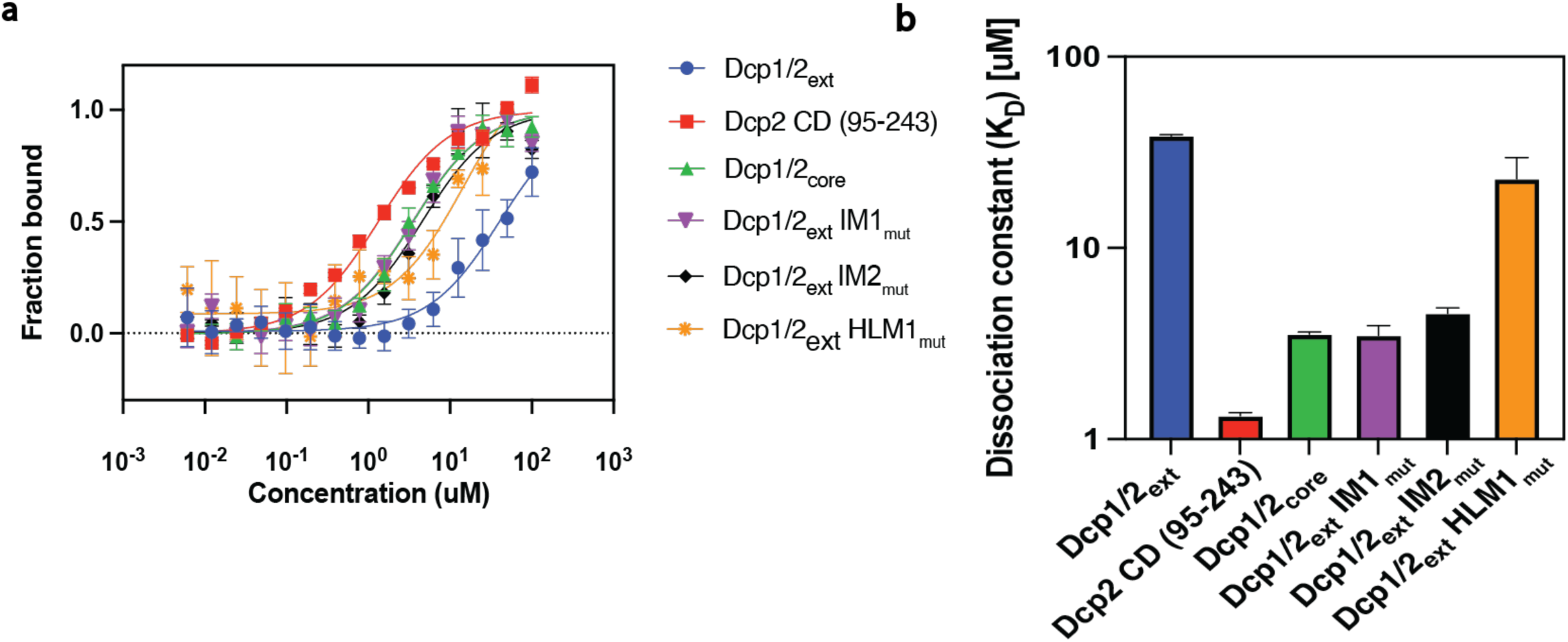
Multiple inhibitory motifs within the Dcp2 IDR repress RNA binding. **a,** Representative binding isotherms showing fraction RNA bound as a function of increasing concentration of Dcp1/Dcp2 complexes. Constructs include Dcp1/Dcp2_ext_ (WT), Dcp2 CD (95–243), Dcp1/Dcp2_core_, and Dcp1/Dcp2_ext_ IDR variants harboring mutations in IM1, IM2, or HLM1. Data represent the mean of three independent experiments; error bars indicate standard deviation. **b,** Dissociation constants (K_D_) derived from fits to the binding curves in panel **a**, plotted on a logarithmic scale.

In contrast, HLM1_mut_ did not significantly alter RNA affinity, as reflected by K_d_ values similar to wild type. This distinction underscores a functional partitioning of sequence motifs within the IDR, with IMs primarily regulating substrate access and HLMs appearing dispensable for RNA binding. Instead, they contribute to multivalent interactions that promote cofactor engagement and higher-order assembly (He and Jacobson 2015; Paquette et al. 2018; Jonas and Izaurralde 2013; Fromm et al. 2012; Harigaya et al. 2010; Tibble et al. 2021).

Together, these findings demonstrate that the Dcp2 IDR modulates substrate recognition through discrete inhibitory motifs that engage in intramolecular interactions.

By selectively tuning access to the RNA-binding channel, these motifs reinforce autoinhibition of the decapping complex and provide a mechanistic link between IDR sequence composition and substrate affinity.

### Mutations to sequence motifs disrupt multivalent interactions in condensates

Previously, we showed that multivalent interactions promoting LLPS of Dcp1/Dcp2 depend on condensate composition (Tibble et al. 2021). In the absence of Edc3, condensate formation required intact IM1 and IM2 and stoichiometric amounts of Dcp1. In contrast, these elements were dispensable for condensate formation of Dcp1 and Dcp2 in the presence of Edc3.

To test whether aromatic motifs in IM1 and IM2 play major roles in Dcp1/Dcp2 condensate formation, we reconstituted Dcp1/Dcp2_ext_ condensates harboring point mutations within the C-terminal IDR, as described above (Fig. 1a). Dcp1/Dcp2 complexes bearing gain-of-function mutations in IMs retained the ability to undergo LLPS in a concentration-dependent manner, indicating that relief of autoinhibition that activates decapping in solution does not abolish the intrinsic capacity of the decapping complex to phase separate (Fig. 3a–d). Notably, IM1_mut_ and IM2_mut_ variants formed visible condensates at concentrations at least twofold below the critical concentration required for wild-type complexes suggesting that disruption of autoinhibitory contacts promotes condensate formation. In contrast, condensates formed by HLM1_mut_ complexes displayed decreased droplet formation and altered material properties, appearing flatter and less liquid-like than those formed by wild-type or IM1_mut_ and IM2_mut_ complexes (Fig. 3d). These morphological differences suggest that HLM1 contributes to LLPS through scaffold-like interactions that shape condensate organization, without being required as a primary driver of droplet formation. As expected from prior structural and biochemical studies showing Edc3 binds HLMs, mutations in IM motifs within the Dcp2 C-terminus did not abrogate Edc3 partitioning into condensates (Fromm et al. 2012). However, IM2_mut_ altered the extent and spatial pattern of Edc3 co-localization within droplets, indicating changes in condensate composition and internal organization (Supplementary Fig. 3). We conclude IMs within the Dcp2 IDR modulate the connectivity and balance of multivalent interactions within condensates. HLM1 emerges as a key regulatory element that contributes to condensate organization by shaping multivalent interaction networks.

**Figure 3.**
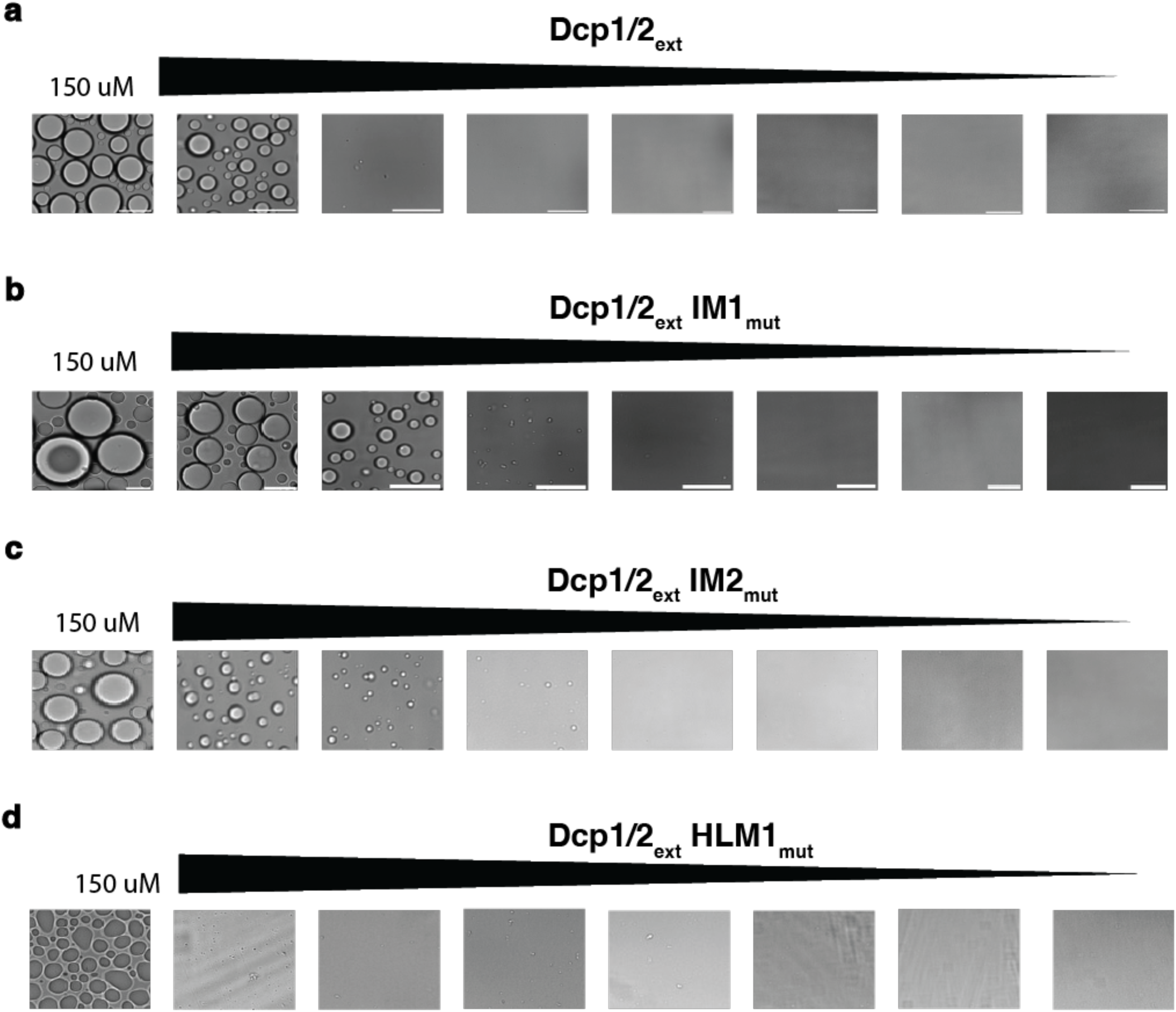
Relief of autoinhibition alters multivalent interactions and condensate assembly. **a–d,** Brightfield microscopy images of Dcp1/Dcp2_ext_ (WT) and Dcp1/Dcp2_ext_ IDR variants harboring mutations in IM1, IM2, or HLM1, showing concentration-dependent phase separation. For each construct, images represent a two-fold serial dilution series (150 µM starting concentration), progressing from left to right. Representative micrographs illustrate differences in droplet formation and persistence across constructs as concentration decreases.

### Lesions in IMs bypass Edc3-dependent activation within condensates

Reconstituted Dcp1/Dcp2 condensates have previously been shown to repress decapping activity in vitro, whereas addition of Edc3 relieves inhibition and strongly activates decapping. These observations suggest that the autoinhibited conformation of Dcp1/Dcp2 adopted in the dilute phase is further stabilized within condensates. We therefore tested whether mutations in aromatic sequence motifs within the Dcp2 IDR that activate decapping in the dilute phase are sufficient to activate decapping within condensates.

To measure decapping in condensates we utilized a dual-labeled 38-mer RNA substrate bearing a 5′ m⁷GDP moiety conjugated to fluorescein isothiocyanate (m⁷GDP–FITC) and a 3′ adenosine coupled to Cy5 (Cy5–RNA body). This substrate enables simultaneous monitoring of cap hydrolysis and RNA retention within the dense phase (Tibble et al. 2021; Depaix et al. 2021). Wild-type Dcp1/Dcp2_ext_ or IDR mutant decapping complexes were incubated with substrate at concentrations exceeding the critical-concentration threshold for phase separation, and reactions were initiated by the addition of MgCl₂. Decapping kinetics were quantified by monitoring loss of m⁷GDP–FITC fluorescence over time.

Within wild-type Dcp1/Dcp2_ext_ condensates, only modest loss of m⁷GDP–FITC fluorescence was observed after 20 minutes, corresponding to an apparent rate constant of ∼0.03 min⁻¹ (Fig. 4a, d). Addition of Edc3 stimulated decapping approximately 14-fold, consistent with its established role as a decapping activator and its enrichment within biological condensates. Importantly, the Cy5-labeled RNA body remained localized within condensates throughout acquisition, indicating that the observed loss of m⁷GDP–FITC signal reflects cap hydrolysis rather than diffusion of RNA out of the dense phase (Fig. 4a, d).

**Figure 4.**
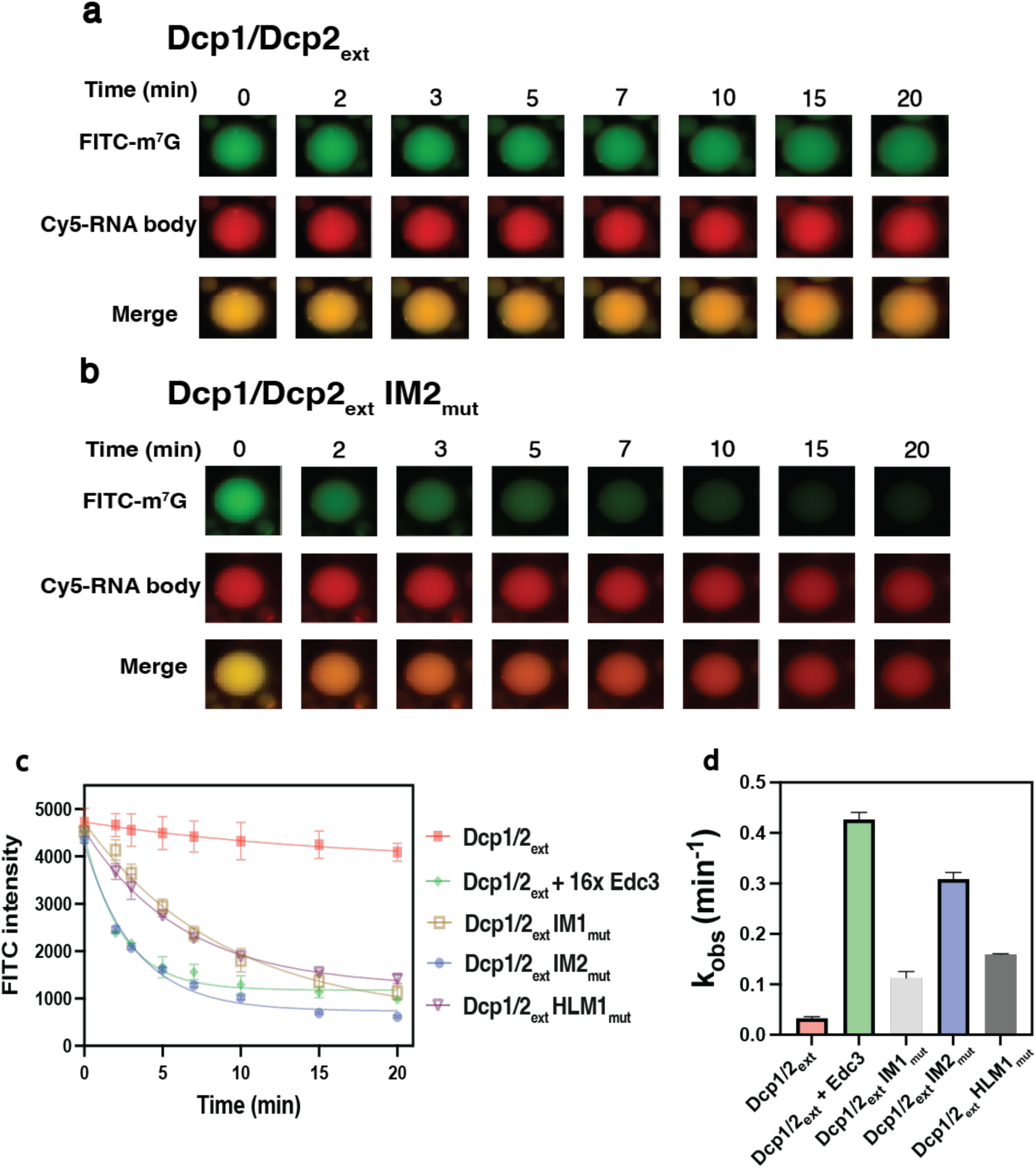
Relief of autoinhibition enhances decapping within reconstituted condensates. **a,** Time-lapse fluorescence microscopy of reconstituted Dcp1/Dcp2_ext_ (WT) condensates (75 µM total protein), monitoring FITC–m^7^GDP (green) and Cy5-labeled RNA body (red) over 20 minutes. WT condensates show minimal time-dependent loss of FITC–m^7^GDP signal. **b,** Time-lapse imaging of condensates formed by Dcp1/ Dcp2_ext_ IM2 (75 µM total protein), demonstrating progressive loss of FITC–m^7^GDP fluorescence over time, consistent with decapping activation. **c,** Quantification of FITC–m^7^GDP fluorescence intensity within condensates over time for the indicated constructs. Data represent mean values from independent experiments; error bars indicate variability between experiments. **d,** Observed rate constants (k_obs_) for decapping within reconstituted condensates, derived from fits to the decay curves shown in panel **c**.

Strikingly, IM2_mut_ resulted in robust decapping activity within reconstituted Dcp1/Dcp2 condensates. IM2_mut_ exhibited an apparent decapping rate constant of ∼0.3 min⁻¹, representing an approximately tenfold increase relative to wild-type condensates (Fig. 4b, d). Notably, this activation occurred in the absence of Edc3, indicating that disruption of IM2-mediated inhibitory interactions is sufficient to convert the condensate environment from a repressive to a catalytically active one. In contrast, IM1_mut_ produced a more modest increase in decapping activity within condensates, yielding an apparent rate of ∼0.1 min⁻¹ (Fig. 4d). The fold-enhancement of decapping activity for IM1_mut_ and IM2_mut_ in condensates differs from that observed in the dilute phase. While the fold enhancement caused by the IM2_mut_ is comparable in dilute and condensed phases, the fold enhancement caused by the IM1_mut_ is less in the condensed phase. This difference, along with the observation that IM1_mut_ decreases the critical concentration of LLPS for Dcp1Dcp2, suggests that IM1 contributes to multivalent interactions that support condensate integrity and internal organization in addition to autoinhibition.

HLM1_mut_ resulted in a modest increase in decapping activity within condensates, consistent with the minimal effects of HLM1 mutation on decapping observed in the dilute phase. Together with the altered material properties of HLM1_mut_ condensates, these findings suggest that HLM1 primarily functions as a scaffolding element that organizes multivalent interactions within condensates, rather than directly regulating the catalytic state of the enzyme (Fig. 3d). We conclude that the inhibitory motifs within the Dcp2 IDR serve dual functions by: 1- stabilizing autoinhibited conformations of the decapping complex and 2- shaping the interaction networks that define condensate organization.

## Discussion

mRNA decapping is a highly regulated step in cytoplasmic mRNA turnover, and the Dcp1/Dcp2 decapping complex exemplifies how IDRs tune enzymatic activity in response to cellular cues. Although prior studies established that the C-terminal IDR of Dcp2 regulates decapping, the lack of residue-level resolution has limited mechanistic insight into how individual sequence motifs influence substrate recognition, catalytic activity, and the conformational landscape of Dcp1/Dcp2. Here, we dissect inhibitory motifs within the Dcp2 IDR and show that conserved aromatic residues within IM1 and IM2 are key determinants of autoinhibition and RNA binding. Our results further demonstrate that these residues contribute to the function and assembly of biomolecular condensates.

Point mutations targeting conserved aromatic residues in IM1 or IM2 robustly activate decapping in the dilute phase, producing effects that phenocopy removal or truncation of the Dcp2 IDR (Fig. 1b, c). These findings extend existing models of Dcp1/Dcp2 regulation by demonstrating that discrete sequence elements within the IDR cooperatively encode autoinhibition. Disruption of these elements is predicted to shift the conformational ensemble of the enzyme toward an active state. The magnitude of activation observed upon mutation of IMs, together with the modest effects of HLM1 mutation alone, supports a model in which autoinhibitory interactions play a dominant role in regulating catalytic activity under single-turnover conditions.

Consistent with this interpretation, IM1_mut_ and IM2_mut_ significantly enhance RNA binding, restoring affinities close to those observed for IDR-truncated or core-only constructs (Fig. 2). These observations support a model in which IMs function to stabilize a closed, inactive conformation of the Dcp1/Dcp2 complex that restricts access to the RNA-binding channel (Wurm et al. 2017; Paquette et al. 2018; Tibble et al. 2021; Deshmukh et al. 2008). In this framework, the Dcp2 IDR forms transient intramolecular contacts with the structured core domains, limiting the complex’s ability to adopt an open, substrate-accessible conformation. Relief of these interactions, either through mutagenesis or through binding of decapping co-activators, promotes RNA engagement and productive catalysis.

IDRs are frequently enriched in phase-separated biomolecular condensates, where they contribute to dense networks of multivalent protein-protein and protein-nucleic acid interactions that drive condensate assembly and organization (Banani et al. 2017; Tibble et al. 2021; Hyman et al. 2014; Brangwynne et al. 2015). Additionally, sequence-encoded molecular grammars within IDRs have been shown to promote composition-dependent LLPS in vivo and in vitro (Rao and Parker 2017; Luo et al. 2018; Wang et al. 2018; Riback et al. 2020). A central finding of this work is that the same sequence motifs that regulate decapping in the dilute phase also contribute to the multivalent interaction network governing condensate formation. Previously, it was shown that deletion of IMs modestly abrogates Dcp1/Dcp2 phase separation (Tibble et al. 2021).

By contrast, using more targeted point mutations of aromatic residues in IM1 and IM2 we were able to uncouple the roles of these regions in condensate formation from their roles in autoinhibition (Fig. 3). Although the IM mutations examined here do not eliminate condensate formation, they alter the internal organization of condensates, particularly the localization of Dcp1/Dcp2 and Edc3 (Supplementary Fig. 3). These results reveal a coupling between autoinhibition and the multivalent interaction network that defines condensate architecture. While HLMs are known to recruit Edc3 directly, our data suggest that IMs also influence these interactions, either by contributing additional interaction valency or by modulating the spatial presentation of binding motifs within the IDR. In this context, HLM1 likely facilitates productive engagement with Edc3 and other cofactors, thereby shaping the condensate environment in which decapping occurs.

Insights from both in vivo and in vitro studies demonstrate that multivalent interactions drive the formation of biomolecular condensates with distinct physicochemical environments (Brangwynne et al. 2009; Li et al. 2012; Banani et al. 2017; Borgia et al. 2018; Tibble et al. 2021). The specific enzymes and nucleic acids enriched within these compartments define their functional identity, generating localized microenvironments that alter reaction kinetics and diffusion in and out of condensates (Ditlev et al. 2018; Peeples and Rosen 2021; Shin and Brangwynne 2017; Stroberg and Schnell 2018; Banani; Su et al. 2016; Prouteau and Loewith 2018). Although condensates are often proposed to enhance enzymatic activity through local concentration effects, accumulating evidence indicates that condensates can also repress activity, depending on their internal architecture and molecular composition (Banani et al. 2017; Peeples and Rosen 2021; Stroberg and Schnell 2018) (Tibble et al. 2021; Riback et al. 2020; Lyon et al. 2021). Previously, reconstituted Dcp1/Dcp2 condensates have been reported to strongly repress decapping activity. Strikingly, IM2_mut_ is sufficient to bypass the requirement for Edc3 and activate decapping within condensates (Fig. 4c, d). These results indicate that relief of autoinhibition is both necessary and sufficient to promote catalytic activity in the condensed phase and suggests that Edc3 primarily functions to activate decapping in condensates by shifting a conformational equilibrium governed by IM2 (Fig. 5a). Our findings help reconcile general expectations that the increased local concentration in a condensate promotes catalytic activity with observations that phase separation of Dcp1/Dcp2 causes inhibition of catalytic activity. Specifically, our observation that mutation of aromatic residues in IMs alleviates autoinhibition and lowers the critical concentration for LLPS suggests that increased phase separation by itself does not repress decapping by Dcp1/Dcp2 in the absence of Edc3 (Tibble et al. 2021). Instead, our data support a more sophisticated model in which enzymatic activity within condensates is controlled by a context-dependent rewiring of the interactions mediated by the Dcp2 IDR (Fig 5). IDR mediated LLPS may entail a change from interactions between the IDR and core domains in Dcp1/Dcp2 in *cis* to intermolecular interactions in *trans* during nucleation on pathway to liquid-liquid phase separation (Fig 5b and c). Together, these findings challenge conventional models of phase separation, which posit that enhanced enzymatic activity arises primarily from local concentration effects. Conformational control of enzyme activity may be a general principle of regulating biochemical reactions within phase-separated compartments such as P-bodies.

**Figure 5.**
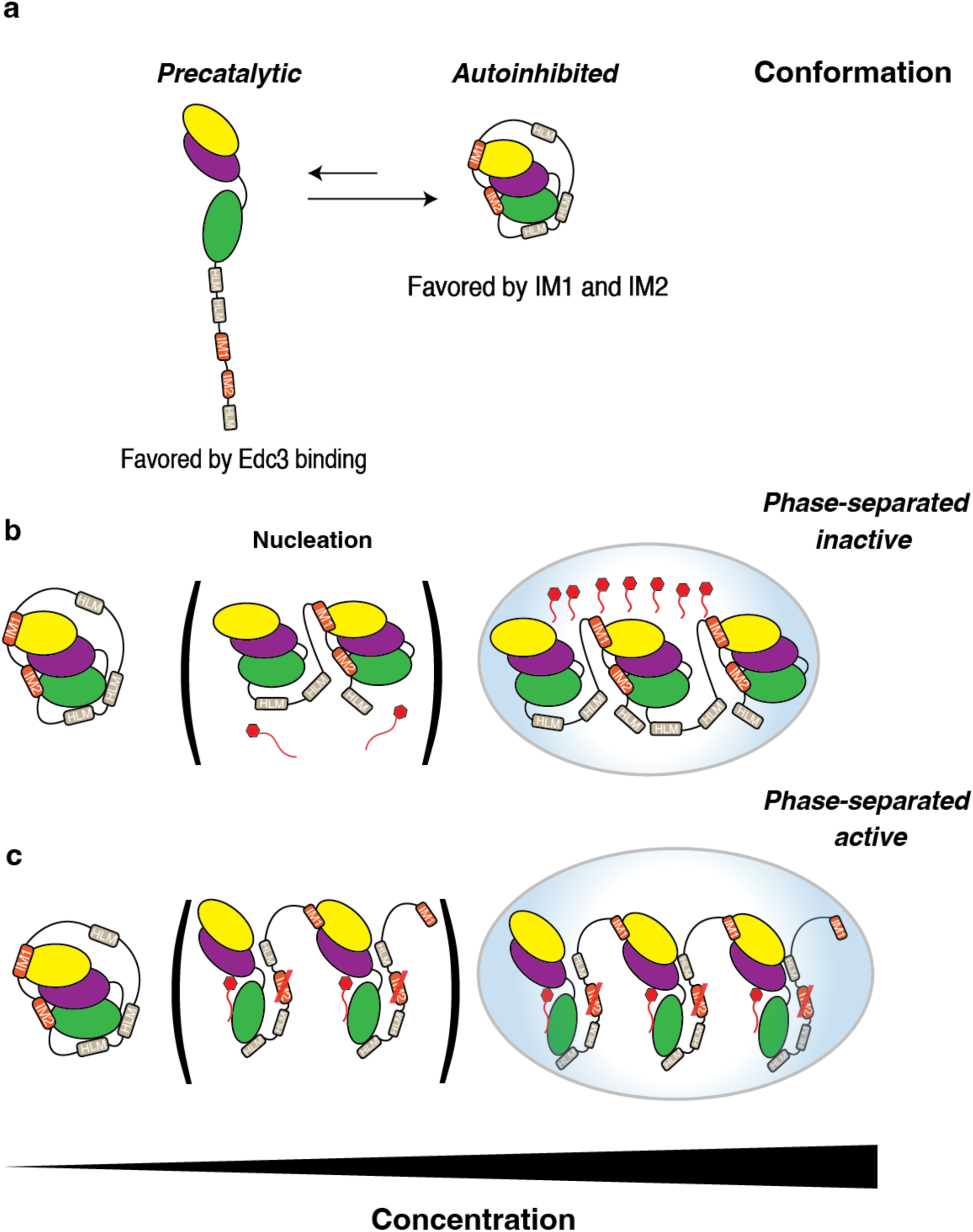
Autoinhibitory interactions within the Dcp2 IDR couple conformational regulation to phase separation and decapping activity. **a,** Left: In solution, Dcp1 (yellow) associates with the Dcp2 N-terminal regulatory domain (NRD, purple) and catalytic domain (CD, green) in a precatalytic conformation that readily engages Edc3 to stimulate decapping. Right: intramolecular contacts involving IM1 and IM2 stabilize a closed, autoinhibited conformation that suppresses decapping activity. **b,** In the wild-type complex, autoinhibitory interactions limit productive RNA engagement and bias Dcp1/Dcp2_ext_ toward an inactive conformation that is further stabilized upon phase separation. **c,** Mutation of IM2 relieves autoinhibition and activates decapping within condensates, while remaining multivalent interactions driven by IM1 promote condensate assembly.

Our work suggests the aromatic residues within IMs of the Dcp2 IDR repress decapping to differing degrees in condensates and the dilute phase. Disruption of either IM1 or IM2 within the Dcp2 IDR is sufficient to relieve autoinhibition in the dilute phase, consistent with a mechanism in which these motifs stabilize the closed conformation through interactions with the structured core. In contrast, relief of autoinhibition in the condensed phase specifically requires disruption of IM2, whereas mutation of IM1 produces only a modest alleviation of repression. These findings suggest a functional partitioning among inhibitory motifs within the Dcp2 IDR. IM2 acts as a key regulatory switch controlling access to the active state in both dilute and condensed phases, whereas IM1 contributes to autoinhibition in the dilute phase but also reinforces intermolecular interactions that organize the multivalent interaction network within condensates (Fig. 5b, c). Thus, sequence elements that enforce autoinhibition in the dilute phase also influence bulk-scale organization of decapping machinery, providing a mechanistic link between intramolecular regulation and the emergent properties of phase-separated compartments. Because prior studies indicate Dcp1 and the IDR of Dcp2 shift a dynamic equilibrium between an open, precatalytic and closed, inactive form of Dcp2 in dilute phase, the mutagenesis studies on IM1 and IM2 presented here suggest liquid-liquid phase separation further represses decapping activity through a conformational selection mechanism akin to the way crystallization can trap one of several specific conformations of enzymes in solution (She et al. 2008; Wurm et al. 2017; Tibble et al. 2021). Rigorously testing this idea requires structural studies of conformational landscape of Dcp1/Dcp2 in condensates, which is a challenge for the future.

AlphaFold3 (AF3) predictions provide additional structural insight into this regulatory mechanism (Abramson et al. 2024). Comparison of the predicted local distance difference test (pLDDT) values between WT and IM2_mut_ models reveals similar levels of predicted disorder in the IDR and structured core domains, indicating that mutation of IM2 does not globally disrupt protein structure (Supplementary Fig. 4). Instead, predicted structural comparisons suggest that IM2_mut_ shifts the enzyme toward an open, precatalytic conformation. In this model, the catalytic domain of Dcp2 is positioned such that the active site remains accessible, resembling previously described open states. In contrast, the WT model adopts a more compact arrangement in which key catalytic residues D47 and W43 are sterically occluded by Y220, consistent with a closed, inactive conformation (She et al. 2006; Mugridge et al. 2016; Wurm et al. 2017; Tibble et al. 2021) (Supplementary Fig. 5). The IM2 mutation is therefore predicted to disrupt intramolecular contacts that stabilize the closed conformation and to favor a structurally open state consistent with relief of autoinhibition.

More broadly, our findings highlight the importance of short linear motifs within IDRs as integrators of biochemical regulation and mesoscale organization. The same residues that mediate intramolecular autoinhibition also contribute to intermolecular interaction networks within condensates, coupling enzymatic regulation to phase behavior through shared sequence grammar. This dual functionality may represent a general principle for enzymes operating within biomolecular condensates, enabling catalytic control to be coordinated with spatial organization. Such a mechanism may allow cells to fine-tune decapping activity across different cellular states and mRNA decay programs. By modulating the engagement of inhibitory motifs through protein–protein interactions or post-translational modifications, cells could regulate both the partitioning of Dcp1/Dcp2 into P-bodies and the functional outcome of those compartments as sites of mRNA storage or decay. Future work examining how additional cofactors, RNA substrates, or signaling inputs interface with these inhibitory motifs will be essential for understanding how RNA decay pathways are regulated in vivo.

Overall, this study establishes a residue-level framework for understanding how the Dcp2 IDR encodes autoinhibition, RNA binding, and condensate function. These findings reveal that activation within condensates can occur independently of coactivators and underscore the central role of IDR sequence architecture in coordinating enzymatic activity and biomolecular organization within the cytoplasm.

## Materials and Methods

### Protein expression and purification

Dcp1/Dcp2_ext_ constructs were expressed in *Escherichia coli* BL21 (DE3) (New England Biolabs) and grown in LB medium. Cells were grown with shaking at 37 °C until the optical density at 600 nm (OD_600)_ reached 0.5-0.7 and then transferred to 4 °C for 30 minutes before induction with 0.75 mM IPTG. Protein was expressed overnight with shaking for 16-18h at 18 °C, centrifuged at 6000 rpm, resuspended in lysis buffer (25 mM HEPES pH 7.5, 400 mM NaCl, 5 mM DTT, 0.1% Triton X-100) supplemented with a protease inhibitor cocktail (Roche), lysed on ice by sonication, and clarified by centrifugation at 14,500 rpm. Clarified lysate was applied to a 5 mL StrepTrap XT (Cytiva) column equilibrated in lysis buffer and washed with 10 column volumes (CV) high salt buffer (25 mM HEPES pH 7.5, 400 mM NaCl, 5 mM DTT) followed by a second wash with low salt buffer (25 mM HEPES pH 7.5, 100 mM NaCl, 2 mM DTT). Elution of strep-tagged protein was performed with 5 CV elution buffer (25 mM HEPES, pH 7.5, 100 mM NaCl, 2 mM DTT, 50 mM biotin). For Edc3 and Dcp1/Dcp2 constructs lacking a C-terminal strep tag, immobilized metal affinity chromatography (IMAC) using a 5 mL HisTrap FF (Cytiva) was used in place of a StrepTrap XT column. For HisTrap purification, the lysis and wash buffers described previously were supplemented with 20 mM imidazole. Proteins were eluted from the HisTrap column with 5 CV low-salt wash buffer containing 300 mM imidazole and incubated with a 1:40 molar ratio of tobacco etch virus (TEV) protease overnight at room temperature. Following elution from StrepTrap XT or TEV digestion, proteins were applied to a HiTrap Heparin (Cytiva) column equilibrated in heparin A wash buffer (25 mM HEPES, pH 7.5, 100 mM NaCl, 2 mM DTT). Protein was eluted over a 12 CV linear gradient to 100% high salt heparin B elution buffer (25 mM HEPES, pH 7.5, 1 M NaCl, 2 mM DTT). Proteins were purified to homogeneity using size-exclusion chromatography with a Superdex 200 16/60 or Superdex 75 column (GE Healthcare), equilibrated in storage buffer (25 mM HEPES, pH 7.5, 150 mM NaCl, 1 mM DTT, 5% glycerol). Protein quality was assessed using SDS-PAGE, quantified by A_280_ on a Nanodrop (ThermoFisher), and flash-frozen in aliquots in liquid nitrogen for storage at -80 °C.

### Fluorescent labeling of proteins and RNAs

Fluorescently labeled Dcp1/Dcp2 and Edc3 constructs were prepared by diluting the proteins in size-exclusion buffer, without glycerol, to 5 mg/ml and 1 mg/ml, respectively. Diluted protein was dialyzed overnight at 4 °C against 25 mM HEPES, pH 7.5, and 150 mM NaCl to remove DTT. Following dialysis, proteins were recovered and incubated with a tenfold molar excess of TCEP to reduce accessible cysteines. Cy5 or Fluorescein maleimide (ThermoFisher) was added at a fivefold molar excess and incubated in the dark at room temperature for 1 hour. The labeling reaction was quenched by adding 2-mercaptoethanol to a final concentration of 10 mM, and the free dye was separated from the labeled protein by size-exclusion chromatography (25 mM HEPES, pH 7.5, 150 mM NaCl, 1 mM DTT). Protein concentration and degree of labeling were determined using UV-Vis spectroscopy on a Nanodrop (ThermoFisher). All fluorescently labeled oligonucleotides were purchased from IDT and contained 5’ phosphorylation and 3’ 6-FAM modifications.

### Fluorescence polarization

Fluorescence polarization assays were performed in Greiner Bio-One 384-well low-volume, non-binding plates. All assays were carried out in 25 mM HEPES, pH 7.5, 100 mM NaCl, 5 mM MgCl_2_, 0.02% Triton X-100, and 0.1 mg/ml acetylated BSA with 10 nM 5’-phosphorylated 30U RNA with 3’ 6-FAM modification (IDT). Reactions were prepared in triplicate and incubated in the dark for 30 min before polarization was measured on the plate reader. A G-factor correction of 1.265 was applied to account for the loss of polarized light through the filter. Equilibrium dissociation constants (*K_d_*) were fit to the equation for one-site-specific binding.

### Single-turnover decapping kinetics

5’-triphosphate 29-mer RNA was enzymatically capped with GTP [α-^32^P] using the Vaccinia virus Capping System (New England Biolabs). Decapping reactions were performed in 25 mM HEPES, pH 7.5, 150 mM NaCl, and 5 mM MgCl_2_ as described previously. Reactions were initiated by mixing 11 μL 2x protein solution with 11 μL 2x RNA [α-^32^P]/MgCl_2_ solution. The final concentration of Dcp1/Dcp2 was 500 nM, and the final concentration of RNA was <10 nM. Time points were taken by quenching in 0.5 M EDTA, spotted onto pre-cut TLC plates (Sigma-Aldrich), and run in 0.75 M LiCl to resolve the accumulated m7GDP product. Quantification was performed using a GE Typhoon scanner and ImageJ software. The fraction m7GDP over time was plotted and fitted to a one-phase association to obtain the rate, *k_obs_.* The rate of decapping, *k_obs_,* reflects subsaturating conditions and is proportional to k_max_/K_m_, where k_max_ is the rate of the catalytic step of decapping and K_m_ is the enzyme concentration that yields a rate of ½ k_max_. For decapping assays where kinetics were too slow to measure using a one-phase association model, rates were determined using simple linear regression.

### Measuring decapping activity within condensates by microscopy

Reconstituted Dcp1/Dcp2_ext_ wild-type or IDR mutant condensates (75 µM) were prepared in 25 mM HEPES, pH 7.5, 150 mM NaCl, 1 mM DTT, 1 mM EDTA, pH 8, and 4 U RNase inhibitor. TEV protease was added in a 1:40 molar ratio (TEV: Dcp1/Dcp2) to initiate phase separation upon cleavage of the MBP solubility tag. Dual-labeled RNA was added to these reactions for a final concentration of 100 nM. Initial images were collected in a PEG-Silane passivated, glass-bottom 384-well plate to observe condensate localization. Decapping reactions were initiated with 5 mM MgCl_2_ and allowed to react for 20 minutes. For all experiments, images were captured in both the FITC and Cy5 channels, and the mean droplet intensity was background-corrected and calculated in ImageJ. Mean intensity was plotted against time and fit to a first-order exponential decay function.

### Brightfield and fluorescence microscopy

Brightfield microscopy images were collected on an inverted widefield Nikon Ti2-E 6D microscope equipped with transmitted-light optics and a scientific CMOS camera, using a Plan Apo 60× air objective. Samples were imaged in 384-well glass-bottom plates (Greiner) that were equilibrated with phase separation buffer (25 mM HEPES, pH 7.5, 150 mM NaCl, 1 mM DTT), PEGylated using 20mg/mL PEG-Silane (Laysan Bio, MPEG-SIL-5000), and blocked with 100 mg/mL BSA before sample addition, as described previously (Keenen et al. 2018). To assess condensate formation across a range of concentrations, Dcp1/Dcp2 complexes were prepared at 150 µM and serially diluted in phase separation buffer immediately before imaging. 25 µM of each dilution was transferred to the imaging chamber and allowed to settle briefly at room temperature before image acquisition. Phase separation was initiated by the addition of 1:40 molar equivalents of TEV: Dcp1/Dcp2 to cleave the N-terminal MBP solubility tag. Dcp1/Dcp2/Edc3 condensates were formed by incubating Dcp1/Dcp2 and Edc3 together before TEV addition. Phase separation and condensate settling were allowed to occur for 30 minutes at room temperature to ensure complete cleavage of the MBP tag. For experiments examining the localization of Dcp1/Dcp2_ext_ and Edc3, 2.5% protein concentration was fluorescently labeled. Enrichment within condensates was estimated as the ratio of the average intensity of 10 droplets to that of the surrounding solution. Images were acquired under identical illumination and exposure settings for all samples to allow comparison across the dilution series. Multiple fields of view were collected for each condition, and representative images are shown. Condensate formation was assessed by the appearance of spherical micron-scale droplets characteristic of liquid–liquid phase separation. Image analysis and scale bar generation were performed in FIJI. Brightfield images were exported and assembled into figure panels without nonlinear processing, with only linear contrast adjustments applied uniformly across the dataset.

### AlphaFold3 modeling and structural analysis

AF3 was used to generate structural models of Dcp1/Dcp2 complexes in the presence and absence of IM2 perturbations. Protein sequences corresponding to Dcp1 and Dcp2 constructs were used as input, and multiple independent predictions were generated for each condition to sample conformational variability. Default AF3 settings were used unless otherwise specified. Model confidence was assessed using the predicted local distance difference test (pLDDT), with analyses focused on high-confidence regions corresponding to folded domains. To evaluate conformational changes, predicted models were compared across conditions and aligned to available crystal structures of Dcp1/Dcp2. Structural superpositions and visualization were performed using UCSF ChimeraX. Changes in the relative positioning of the Dcp2 catalytic domain and accessibility of the RNA-binding channel were assessed qualitatively from aligned models. Figures were generated in ChimeraX, with domain organization and key residues annotated based on the predicted structures.

## Acknowledgements

We thank the Center for Advanced Light Microscopy at UCSF for technical assistance with data collection. We thank Gross Lab members Xi Liu for guidance on experimental design and Gabriel Braun for preparing cap-radiolabeled RNA. We also thank Ryan Tibble and Geeta Narlikar for valuable scientific discussions and feedback on the manuscript. This work was supported by the U.S. National Institute of Health (R01 GM148881 to J.D.G.). Molecular graphics and analyses performed with UCSF ChimeraX, developed by the Resource for Biocomputing, Visualization, and Informatics at the University of California, San Francisco, with support from National Institutes of Health R01-GM129325 and the Office of Cyber Infrastructure and Computational Biology, National Institute of Allergy and Infectious Diseases.

## Author contributions

T.A. and J.D.G. designed experiments, and K.G., J.K., and J.J. designed and synthesized the dual-labeled fluorescent RNA probe. T.A. purified proteins and performed all experiments. T.A. and J.D.G. contributed to the editing of the manuscript.

J.D.G supervised all research.

